# The evolution of irreversible cell differentiation under cell death effect

**DOI:** 10.1101/2024.11.25.625286

**Authors:** Yuanxiao Gao, Xueyan Zhao, Caixia Li

**Author notes:** Corresponding author: Yuanxiao Gao.

## Abstract

Cell differentiation is an important characteristic of multicellular organisms which produce new-typed cells to engage in diverse life functions. Irreversible differentiation, as an important differentiation, describes cells differentiated by determined trajectories to form specialized cell types. It has been found that differentiated cell types often show different death rates. Yet, it is still unclear what role cell death plays in shaping the formation of irreversible cell differentiation. Here, we establish a theoretical model to investigate the impact of cell death on the evolution of irreversible cell differentiation in multicellular organisms. Irreversible differentiation refers to the loss of a cell type’s differentiation potential, and it is constructed by the sequences of differentiation probabilities of a cell type across cell divisions. We show that irreversible differentiation is more likely to occur when cell death rates between cell types are linear. Meanwhile, differences in death rates between cell types affect the emergence conditions of irreversible differentiation, whereas no significant impacts on that from equal cell death rates. Additionally, we found that cell death impacts the cell number and cell composition of a mature organism. These findings provide insights into understanding the role of cell death in the formation of cells’ irreversible differentiation.

## 1 Introduction

Cell differentiation, a hallmark of multicellular organisms, accounts for a series of dynamic behaviors and significant morphological changes in the formation of multicellular organisms [Maynard Smith and Szathmáry, 1995, Szathmáry and Smith, 1995, Grosberg and Strathmann, 2007, Mikhailov et al., 2009, Márquez-Zacarías et al., 2021]. Cell differentiation allows complex multicellular organisms to maintain varying life functions through different specialized cell types [Carroll, 2001, McCarthy and Enquist, 2005, Arendt, 2008]. Many mechanisms have been introduced to explain cellular differentiation [Extavour and Akam, 2003, Mikhailov et al., 2009, West and Cooper, 2016, Arendt et al., 2016, Brunet and King, 2017, Márquez-Zacarías et al., 2021] as the emergence of new cell type is still obscure [Arendt, 2008]. Along with newly formed cell types, the cell differentiation process shows different differentiation patterns in terms of the differentiation potential, such as irreversible differentiation and reversible differentiation [Rodrigues et al., 2012, Gao et al., 2021, Nakamura et al., 2024]. The most striking example is germ-soma differentiation which leads to germ-line cells and somatic cells. The two cell types are both the result of an irreversible differentiation process where cells eventually lose their cell differentiation capability in an organism [Matt and Umen, 2016]. However, their cell fates are different as germ-line cells can pass to the next generation and develop into a new organism whereas non-reproductive somatic cells have limited lifetime and eventually undergo senescence and cell death. Thus, cell differentiation produces different types of cells that undergo varying life spans in an organism’s development, depending on cell types.

Cell death is a complex biological process that includes many different typed forms via different mechanisms [Vaux and Korsmeyer, 1999, Fuchs and Steller, 2011, Galluzzi et al., 2018, Bertheloot et al., 2021, Park et al., 2023]. Cell death in multicellular organisms not only occurs at the endpoint of whole biological organisms but throughout an individual’s developmental processes. Even without external environmental stimuli, cells can still undergo cell death throughout organisms’ development known as programmed cell death. Cell death has various functions, including regulation of cell number, elimination of abnormal and dangerous cells, sculpting or deletion of structures [Vaux and Korsmeyer, 1999, Fuchs and Steller, 2011, Park et al., 2023] It has been found that programmed cell death occurs for specific cell types. For example, in *C. elegans*, the programmed death of somatic cells is strictly controlled by cell lineage [Ellis et al., 1986]. Meanwhile, abnormal regulation of programmed cell death in humans accounts for a wide range of diseases, including developmental disorders and cancer [Fuchs and Steller, 2011]. Cell differentiation and cell death, together with cell proliferation serve a crucial relationship in individual’s development and tissue homeostasis [Park et al., 2023]. Cell death is often a consequence of mistakes that occur among cell division and differentiation [Fuchs and Steller, 2011]. In turn, cell death can secrete factors that stimulate cell proliferation and differentiation [Fuchs and Steller, 2011].

Although cell differentiation and cell death have both been extensively investigated, few works have examined their relationship from an evolutionary perspective. Theoretical work of cell differentiation is mostly focusing on finding the optimal distribution of cell types under the division of labour hypothesis of the origin of cell differentiation [Michod, 2007, Willensdorfer, 2009, Gavrilets, 2010, Rossetti et al., 2010, Rueffler et al., 2012, Ispolatov et al., 2012, Solari et al., 2013, Goldsby et al., 2014, Cooper and West, 2018, Liu et al., 2021, Cooper et al., 2021, Gao et al., 2021, Cooper et al., 2022]. Theoretical work of irreversible patterns of cell differentiation has not been paid enough attention previously probably due to lacking a mathematical tool to describe different patterns together with the complexity and high dimensions in such models. Recently, irreversible differentiation has been investigated in multicellular organism models with two cell types via sequences of stochastic cell differentiation probabilities over cell divisions [Gao et al., 2021, 2023]. Irreversibility of cell differentiation was investigated in a model by considering the gene regulatory network [Nakamura et al., 2024]. However, cell death has not been considered in these models though previous studies have shown that natural selection favors cell death in clustered multicellularity as it can promote organisms’ reproduction [Ratcliff et al., 2012, Libby et al., 2014]. Meanwhile, research has also shown that dying cells can affect the cellular environment and trigger tissue regeneration [Fuchs and Steller, 2015]. Thus, it will greatly help researchers to understand the relationship between cell death and differentiation patterns by investigating cell death and its impact on the evolution of irreversible differentiation patterns.

In the work, we propose a theoretical model to study the impact of cell death on the evolution of irreversible cell differentiation based on previous theoretical research [Gao et al., 2023]. Here, the cell differentiation probability can change depending on its division state [Gao et al., 2023]. Cell differentiation patterns are constructed by the sequences of differentiation probabilities across cell divisions. Irreversible differentiation refers to the differentiation probability sequences of a cell type converges to 0. We considered two functionally different cell types: germ-like and soma-like. Cell death is accompanied by cell division and differentiation in an organism. We adopt an organism’s reproductive rate as a proxy of an organism’s fitness, i.e. the expected number of offspring of the organism. The research aims to seek the incipient evolution conditions of irreversible differentiation under varying degrees of cell death in both cell types. Our conclusions show that irreversible differentiation emerges mostly when cell death rates between cell types are linear. The emergence conditions of irreversible differentiation change with different cell death rates but have no significant change under the same cell death rates. Finally, we found that cell death affects the cell number and cell composition of mature organisms.

## 2 Model

In the model, we consider multicellular organisms that can have two cell types and can adopt all potential differentiation strategies (Fig1). Based on previous work where the cell type setting is inspired by *Volovx*, two functionally different cell types are considered: germ-like and soma-like [Gao et al., 2021, 2023]. Germ-like cells are responsible for reproduction whereas soma-like cells are responsible for survival. We should note that the two cell types are the transient cell types rather than the final determined cell types. Thus cells can differentiate to another cell type via cell division. Each organism is born with a germ-like cell and then undergoes division with differentiation and death. Then each cell undergoes synchronous division producing two daughter cells until the organism matures after *n*th cell divisions which is a given value of interest. After maturity, all soma-like cells die and germ-like cells turn into new offspring organisms and repeat the life cycle. During cells’ division, they can differentiate with certain probabilities. For instance, there are three division pathways for germ-like cells: divide into two germ-like cells with a probability of *g*_*gg*_, differentiation into one germ-like cell, and divide one soma-like cell with a probability of *g*_*gs*_, differentiation into two soma-like cells with a probability of *g*_*ss*_. Here, *g*_*gg*_ + *g*_*gs*_ + *g*_*ss*_ = 1. Differentiation probabilities can also be expressed as 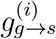 which is 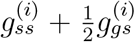, where *i* represents the *i*th cell division. The probabilities between two adjacent cell divisions are randomly taken but with a difference i.e. 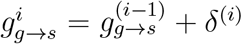, where |*δ*^(*i*)^ ≤ 1|. Soma-like cells divide similarly to germ-like cells.

**Figure 1.**
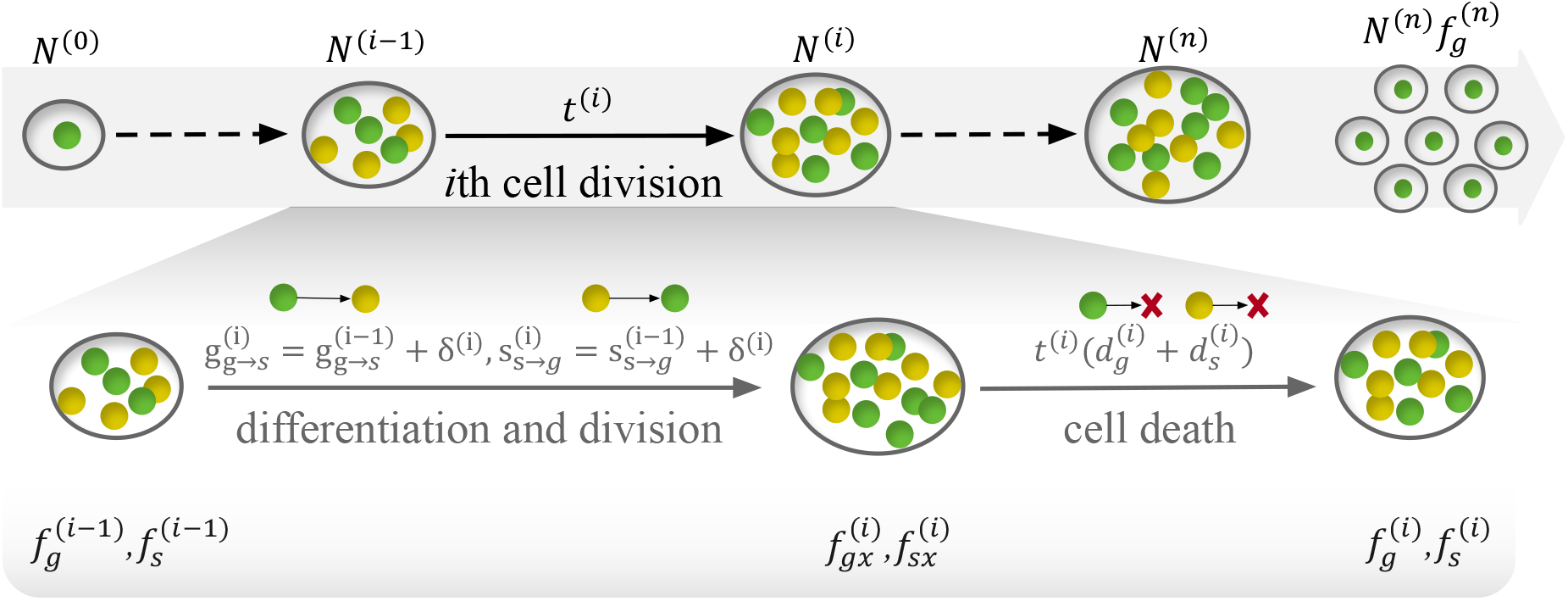
Schematic of the life cycle of an organism with cell death. The green spheres represent germ-like cells, and the yellow spheres represent soma-like cells. An organism is born with a germ-like cell, and cells do synchronous cell divisions with differentiation and cell death until *n* rounds of cell divisions. Then, the germ-like cells are released as offspring. We assume that cells die instantaneously after cell divisions and differentiation. *N*^(*i*)^ is the cell number after the *i*th cell division.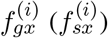 is the proportion of germ-like cells (soma-like cells) after the *i*th cell differentiation 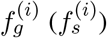 is the proportion of germ-like cells (soma-like cells) after the *i*th cell death. *t*^(*i*)^ is the waiting time from the (*i* − 1)th cell division to the *i*th cell division, see the main text for more detail.

Differentiation strategy is defined by the sets of sequences of differentiation probabilities of cell types. In the sequence, the first element consists of six probabilities 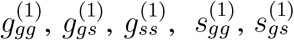, and 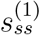 which are randomly chosen to execute cell differentiation. Then, the cell differentiation probabilities for the second cell division are deviated at most *δ*^(1)^. Similar methods for taking the following differentiation probabilities until the organism reaches maturity. We take the same definition of cell differentiation strategy as Gao’s work [Gao et al., 2023]. Cell differentiation patterns include non-differentiation (*ND*), reversible differentiation (*RD*), and irreversible differentiation (*ID*) based on the differentiation probabilities at the last cell division. Among them, non-differentiation (*ND*) means that cells do not differentiate and only proliferate germ-like cells, i.e. 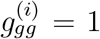 for *i* = 1, 2, …, *n*. Irreversible differentiation (*ID*) indicates the presence of germ-like cell differentiation in the first *n* − 1 divisions i.e. there exists at least once 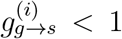 for *i* = 1, …, (*n* − 1). At the last cell division i.e. *i* = *n*, either 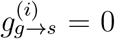 or 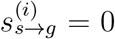. Reversible differentiation (*RD*) is the rest strategy that with 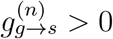 and 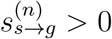.

We measure the performance of differentiation strategies by using an organism’s expected reproductive rate which is the number of germ-like cells after *n*th cell division [Gao et al., 2023]. The cell differentiation strategy which leads to an organism growing fastest will be selected and be denoted as optimal one. The reproductive rate of an organism is defined as:

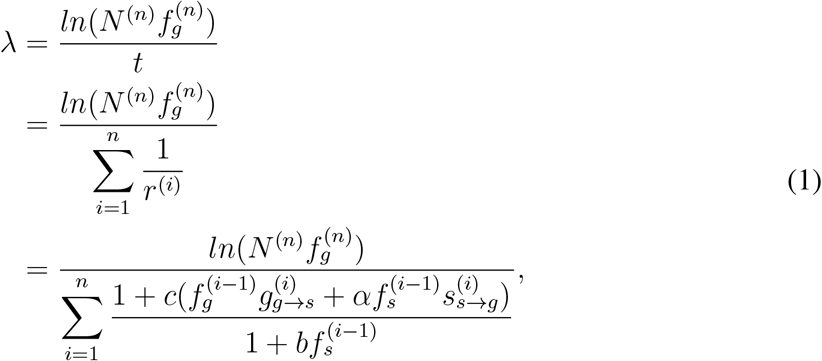

where *N* ^(*n*)^ is the total cell number and 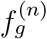 is the fraction of germ-like cells of the mature organism, respectively. *r*^(*i*)^ the cell division rate between the (*i* − 1)th and the *i*th division, which is defined as 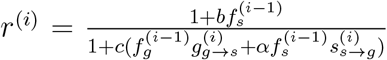. 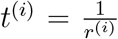 is the expected growth time between the (*i* − 1)th and the *i*th division. Thus, 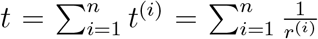 is the expected growth time of an organism. *b, c* and *α* measure the effects of cell interactions. Specifically, *b* and *c* represent the benefits and costs that a differentiation strategy brings to an organism respectively. 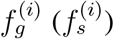 represents the fraction of the germ-like (soma-like) cells after the *i*th cell division.

Next, to calculate an organism’s reproductive rate, we calculate 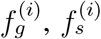, *andN* ^(*n*)^. We assume that 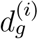 and 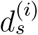 are the death rates per unit time of germ-like cells and soma-like cells at *i*th cell division respectively i.e. during the time between the (*i* − 1)th and the *i*th division. Here, the cell death rate is proportional to the waiting time between adjacent cell divisions (Fig1). We assume that cells die instantaneously after cell divisions and differentiation as cell death is often a consequence of the mistakes that occur among cell division and differentiation [Fuchs and Steller, 2011]. Therefore, we first calculate the change in cell fractions caused by cell division and differentiation, and then cell death. For convenience, we use 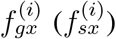 to denote the cell fraction of germ-like cell (soma-like cell) after the *i*th cell differentiation, and 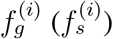 to denote the cell fraction that after the *i*th cell death, thus we have

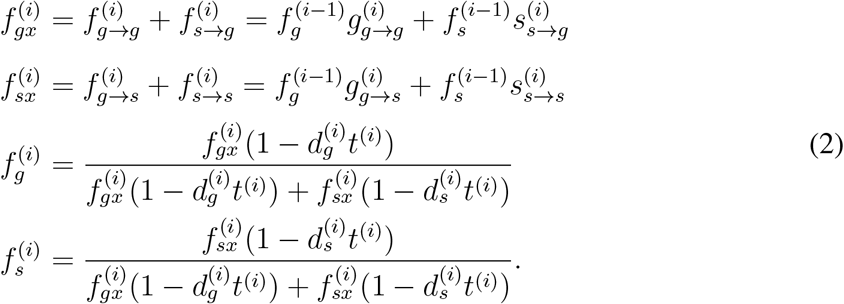

where 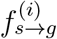 is the fraction of soma-like cells that differentiate into the germ-like cell. *t*^(*i*)^ is the expected time between (*i* − 1)th and *i*th cell division, which is 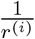. 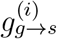 refers to the probability of germ-like cells differentiating into soma-like cells during the *i*th division.

Next, we calculate the total cell number *N* ^(*n*)^. With cell death, the maturity organism size *N* ^(*n*)^ is smaller than 2^*n*^. Specifically, when the unit death rates of germ-like cells and soma-like cells are the same 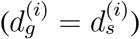, the total number of mature cells *N*^(*n*)^ is

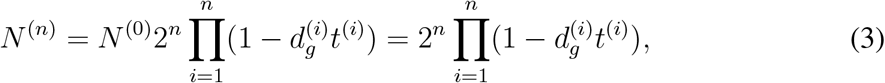

where 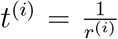. *N*^(0)^ refers to the number of cells for the newborn organism, where there is only one germ-like cell, so *N*^(0)^ = 1. When the unit mortality rates of germ-like cells and soma-like cells are different 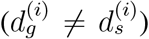, the total number of cells *N*^(*n*)^ at the time of cell development and maturation is

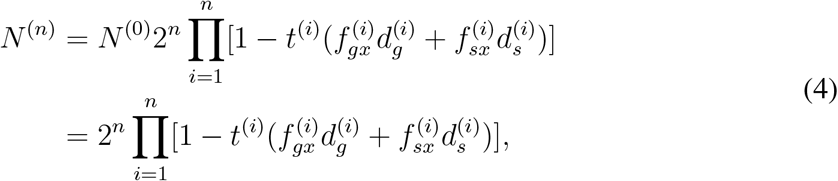

see Appendix A.1 for more detail.

## 3 Results

To investigate the emergence conditions of irreversible differentiation, we numerically calculate the reproductive rates of organisms that adopt all potential differentiation strategies under varying parameters of cell death rates and differentiation benefits and costs, see Appendix A.2 for more detail. Different strategies compete by the organisms’ reproductive rates that the strategies acted on. Then we seek the optimal strategy which has the largest reproductive rate through all parameter space. The parameter space where *ID* is optimal refers to the emergence conditions of irreversible differentiation. For simplicity, we assume that the cell death rate is stage-independent which means 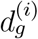 and 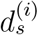 are constant to *i*. We denote them by *d*_*g*_ and *d*_*s*_ in the following investigation. For convenience, we set *δ*^(*i*)^ = 0 or *δ*^(*i*)^ = 0.1. The numerical calculation of the reproductive rate takes the same method as that in the previous study [Gao et al., 2023].

### 3.1 Irreversible differentiation occurs mostly under linear death rates of different typed cells

A high cell death rate will lead to organismal death where no cells can survive to maturity. Therefore, we need to investigate the impact of cell death on organismal death before irreversible differentiation emerges. In the model, the cell death rates of germ-like cells and soma-like cells are *d*_*g*_ and *d*_*s*_ respectively. As each organism is born with a germ-like cell, thus high *d*_*g*_ will lead to the death of germ-like cells which further leads to organismal death. Thus, organisms cannot survive at high *d*_*g*_. Our numerical results show that when the cell death rate of germ-like cells increases to around 1.8, organisms die (Fig 2A). In comparison, a high soma-like cell death rate *d*_*s*_ alone will not lead to organismal death. Under this scenario, organisms that produce few soma-like cells will not die. An extreme example is the organisms with *ND* strategy, where the organism only contains germ-like cells, thus high *d*_*s*_ has no impact. Finally, a combined cell death rate of both cell types will lead to fast organismal death. We found that organisms is likely to die when *d*_*g*_ exceeds 0.5 and *d*_*s*_ exceeds 0.4 (Fig 2A). In the following investigation of differentiation strategies, we only consider the range of cell death rates that ensures organisms’ survival.

**Figure 2.**
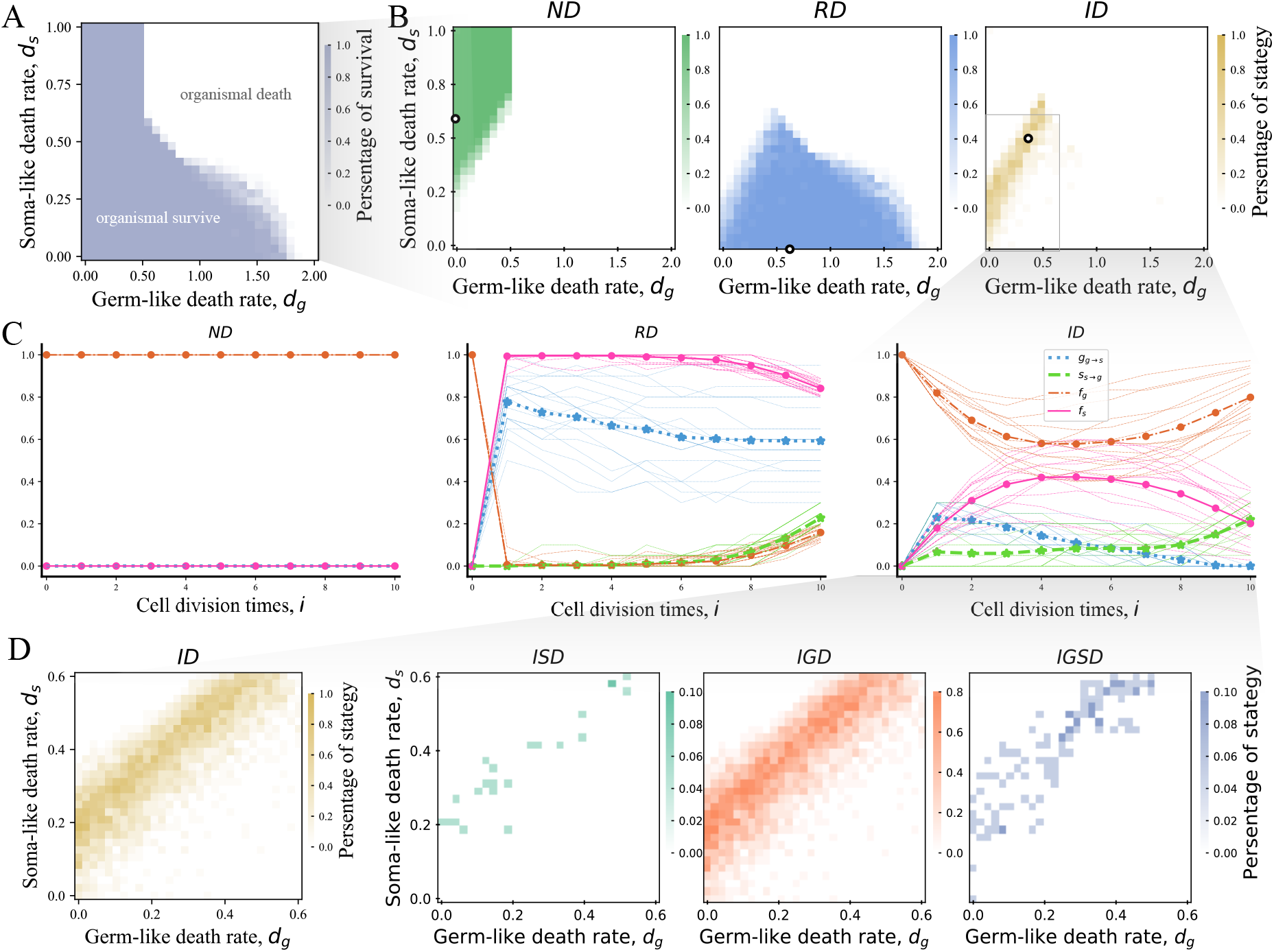
Optimal strategy under cell death rates. **A**.The survival space of organisms under cell death rates. The blue area represents the cell death rate space where cells can survive to mature. The white area indicates that organisms die by losing all their cells. **B**. The proportion of optimal strategy being *ND, RD* and *ID*, respectively. **C**. Cell differentiation probabilities (*g*_*g*→*s*_, *s*_*s*→*g*_) and the frequencies of cells (*f*_*g*_, *f*_*s*_) across cell divisions for the strategies that denoted by the white circles in panel *B*. **D**. The proportion of each optimal sub-strategy of *ID* under cell death rates, respectively. Parameters of all panels: maximal cell division times *n* = 10, 0 *≤ δ*^(*i*)^ *≤* 1, and *b* = *c* = *α* = 1. At each pixel, 20 replicates for calculating the optimal strategy.

Our results show that irreversible differentiation strategy *ID* occurs in the parameter space between the space for strategy *ND* and *RD* emergence (Fig 2B). We found that *ID* evolves in low cell death rate both for germ-like cells *d*_*g*_ and soma-like cells *d*_*s*_. The growth of organisms with *ND* strategy is the clonal reproduction of germ-like cells and no soma-like cells (Fig 2C). Thus, *d*_*s*_ has no impact on *ND. ND* strategy can evolve in high *d*_*s*_. According to formula (3), an organism can survive when its final cell number *N* ≥ 1. That is

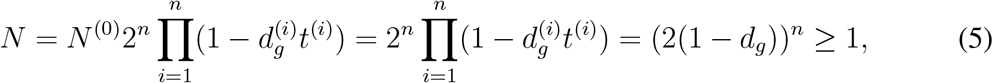

where *t*^(*i*)^ equals 1 in *ND* according to formula (1). Then, organisms that adopted strategy *ND* can survive when *d*_*g*_ ≤ 0.5. Our numerical results are consistent with this conclusion, see Fig 2A and B. Strategies of *RD* evolve at high *d*_*g*_ as the strategy can keep relatively fewer germ-like cells in an organism to avoid cell death. In particular, the optimal strategy of *RD* is the one with a high differentiation probability from germ-like to soma-like and a low differentiation probability from soma-like to germ-like until the few last round of cell divisions, see Fig 2C. The optimal strategy has the opposite cell differentiation probability tendency in the last few cell divisions as soma-like cells die at the end of the multicellular life cycle. Thus, the strategy of *RD* that leads less germ-like cells to avoid cell death but produces as many germ-like cells at maturity to increase the number of offspring is selected as it increases an organism’s reproductive rate. Since the fraction of germ-like cells produced by *ID* can be between that by strategy *ND* and strategy *RD, ID* emerges in the parameter space between the space that emerges for *ND* and *RD* (Fig 2C). Therefore, *ID* is optimal under selection of both *d*_*g*_ and *d*_*s*_.

We further found that *ID* emerges mostly when *d*_*g*_ and *d*_*s*_ fall in a narrow strip space showing a linear relationship, see Fig 2B and D. Even though *ID* can emerge in these narrow strip areas where *d*_*g*_ and *d*_*s*_ do not necessarily show a strictly linear relationship. Nevertheless, the linear condition leads to the highest percentages of *ID* (Fig 2B and D). Notably, the conclusion does not exhibit a significant change when differentiation benefits *b* and costs *c* vary, see Appendix A.3. This conclusion indicates that the two-typed cells need to keep proportional changes in cell death rate for the emergence of irreversible differentiation. The reason is that if there is a big difference between the two cell death rates, then either *ND* or *RD* will be selected. Specially, if the increase of *d*_*g*_ is greater than that of *d*_*s*_, then the strategy *RD* will be chosen. Since *RD* includes the strategy that produces a high fraction of soma-like cells in the early stages of cell divisions and then produces as many germ-like cells as offspring in the late cell divisions. In contrast, if the increase of *d*_*s*_ is greater than that of *d*_*g*_, the strategy *ND* will be chosen as it only contains germ-like cells and thus can avoid cell death.

Furthermore, we investigate what kind of strategy is chosen in *ID*. Based on the definition, we know that at least one cell type has no cell differentiation in *ID*, Thus, *ID* includes three sub-strategies *IGD, ISD*, and *IGSD*, which are according to 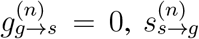 and 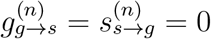 respectively. The three sub-strategies together make strategy *ID* impacted by *d*_*g*_ and *d*_*s*_ simultaneously. We found that *IGD* is the most occurring sub-strategy in irreversible differentiation, see Fig 2D. The finding is consistent with previous study [Gao et al., 2023].

### 3.2 Cell death rate differences between different cell types affect the emergence condition of irreversible differentiation

In this section, we investigate the impacts of cell death on the emergence condition of irreversible differentiation under differentiation benefits *b* and costs *c*. An organism will grow fast by producing soma-like cells which serve survival functions. Differentiation benefits measure the contribution of soma-like cells in an organism. The contribution will increase cell division rates and thus decrease an organism’s growth time [Gao et al., 2023]. Differentiation costs inhibit cell differentiation and punish cell differentiation by decreasing an organism’s reproductive rate, see formula (1). We found that different cell death rates largely change the evolving conditions of irreversible differentiation, whereas it is unchanged under equal cell death rates, see Fig 3.

**Figure 3.**
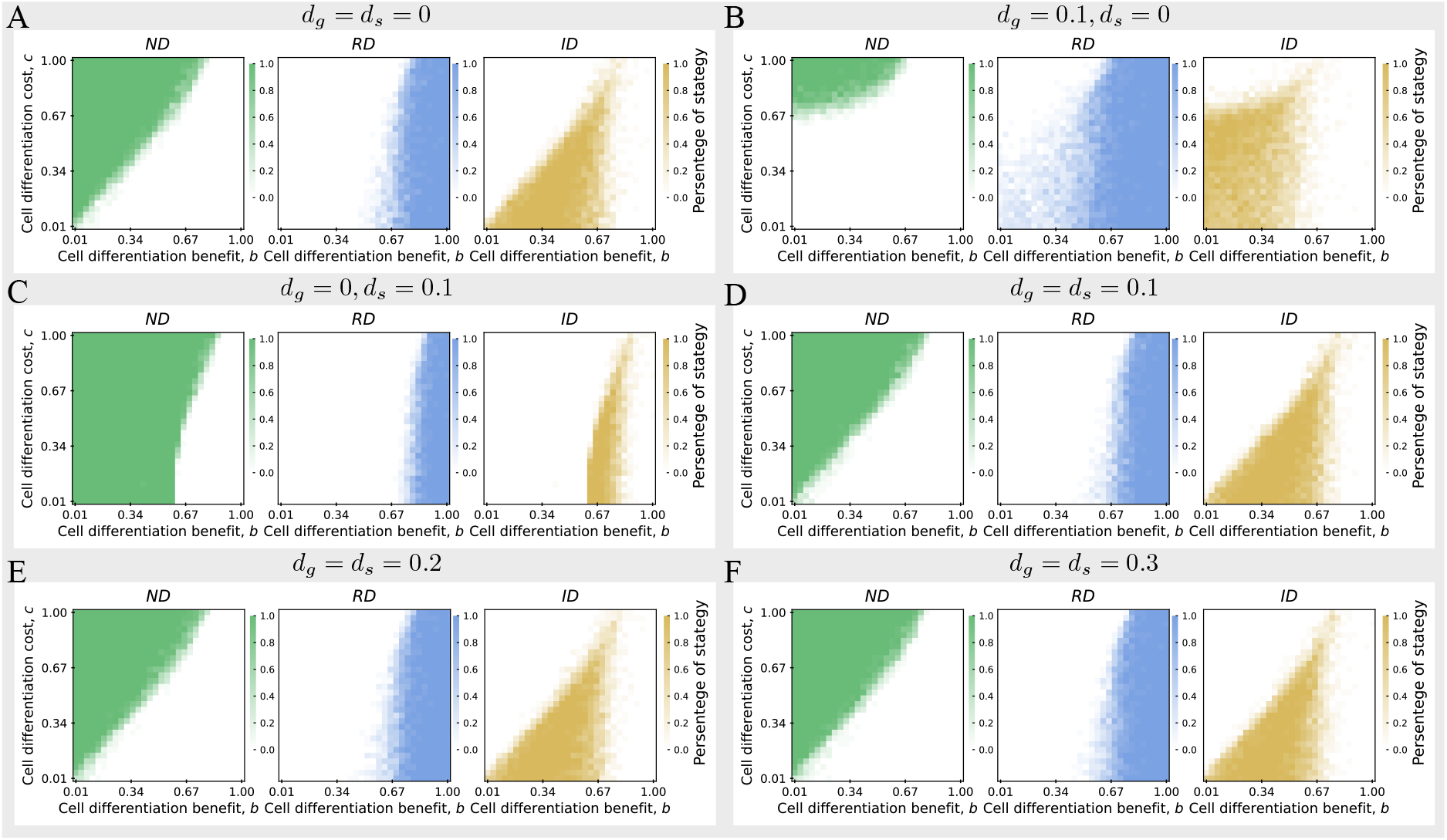
The percentage of optimal differentiation strategy under differentiation benefits and costs with varying cell death rates. The percentage of optimal differentiation strategy being *ND, RD*, and *ID* under differentiation benefits and differentiation costs in the condition of 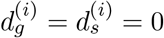 (panel **A**), 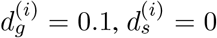 (panel **B**), 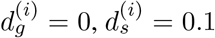 (panel **C**), 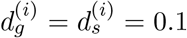 (panel **D**),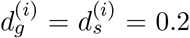 (panel **E**), 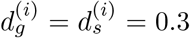 (panel **F**). When the cell death rates of germ-like cells and soma-like cells are equal, although the cell death rate increases, its impact on the optimal strategy is not significant. Other parameter: *n* = 10, 0 ≤ *δ*^(*i*)^ ≤ 1, *α* = 1. At each pixel, 20 replicates for calculating the optimal strategy.

Without cell death, high differentiation costs prohibit cell differentiation, thus *ND* emerges at the area of high *c* and low *b*, see Fig 3A. Comparatively, when the differentiation benefits are enough large, differentiation strategy *RD* will be the optimal strategy. *ID* emerges in the parameter space with a certain amount of differentiation benefits and low differentiation costs. The conclusion is consistent with previous work [Gao et al., 2021, 2023]. When germ-like cell death rate increases, differentiation strategies of *ID* and *RD* replace *ND* being the optimal one in the area with low differentiation benefits and low differentiation costs, see Fig 3B. This is due to the fact that organisms with *ND* only have germ-like cells and thus have more cell death compared with organisms that adopted the other two strategies under the scenario of *d*_*g*_ but without *d*_*s*_. In contrast, when soma-like cell death increases, *ND* will invade the parameter space which is occupied by *ID* and *RD*. As differentiation strategies (*RD* and *ID*) contain soma-like cells, they are more sensitive to soma-like cell death. Our results show that the evolving conditions of *ID* change differently by increasing the same amount of death rate on the two cell types. *ID* is likely to be found in the organism with germ-like cell death rather than soma-like cell death (Fig 3B and C).

However, most interestingly, we found that the conditions for evolving *ID* are unchanged much when both cell types possess the same death rate, Fig 3A, D-F. The same death rate for both cell types will equally decrease the number of both cells but does not change their proportions. Thus, the same death rate impacts organisms in the same way and eventually reduces an organism’s cell number at the maturity stage. However, it does not impact the evolving condition for different differentiation strategies. The result indicates that the same cell death rate for different cell types will not change the emergence conditions of the differentiation patterns. Therefore, for the evolution of more complex and hierarchical structures in an organism, cells need to have different cell death rates. This finding indicates that cell death rate plays an important role in the formation of cells’ differentiation patterns.

### 3.3 Cell death impacts organisms’ size and cell composition

Finally, we investigate the impact of cell death on organismal size and cell composition in mature organisms which adopted *ID* as the optimal strategy. Here, the organismal size is the total cell number that an organism can reach after *n* rounds of cell divisions. The cell composition is investigated by the fraction of germ-like cells. We found that organismal size decreases with both cell death rates *d*_*g*_ and *d*_*s*_, see Figure 4. Meanwhile, the proportion of a cell type is inversely proportional to the cell death rate of this type. Without cell death, cells undergo binary synchronous division, and an organism will eventually have 2^*n*^ cells. Therefore, we investigate the changes in organismal size and cell composition for the organism adopted *ID* by comparing them to those that are without cell death. Our numerical results show that the organismal size can be smaller to 1 under both *d*_*g*_ and *d*_*s*_ compared with 1024 cells without cell death for *n* = 10 (Figure 4C). Without cell death, the proportion of germ-like cells is around 79% which means 21% soma-like cells, see Figure 4C. When *d*_*g*_ increases to 0.1, the maturity organismal size decreases from 1024 to 723, and the proportion of germ-like cells decreases from 0.79 to 0.42. However, when *d*_*s*_ increases to 0.1, the maturity organismal size decreases from 1024 to 879, and the proportion of germ-like cells increases from 0.79 to 0.92. Comparatively, the amount of organism size decreases more than two folds for *d*_*g*_ decrease than that for *d*_*s*_ decrease, and the amount of the fraction of germ-like cells decreases more than three folds. These results demonstrate that the death of germ-like cells plays a more crucial role in influencing cell number and cell composition.

**Figure 4.**
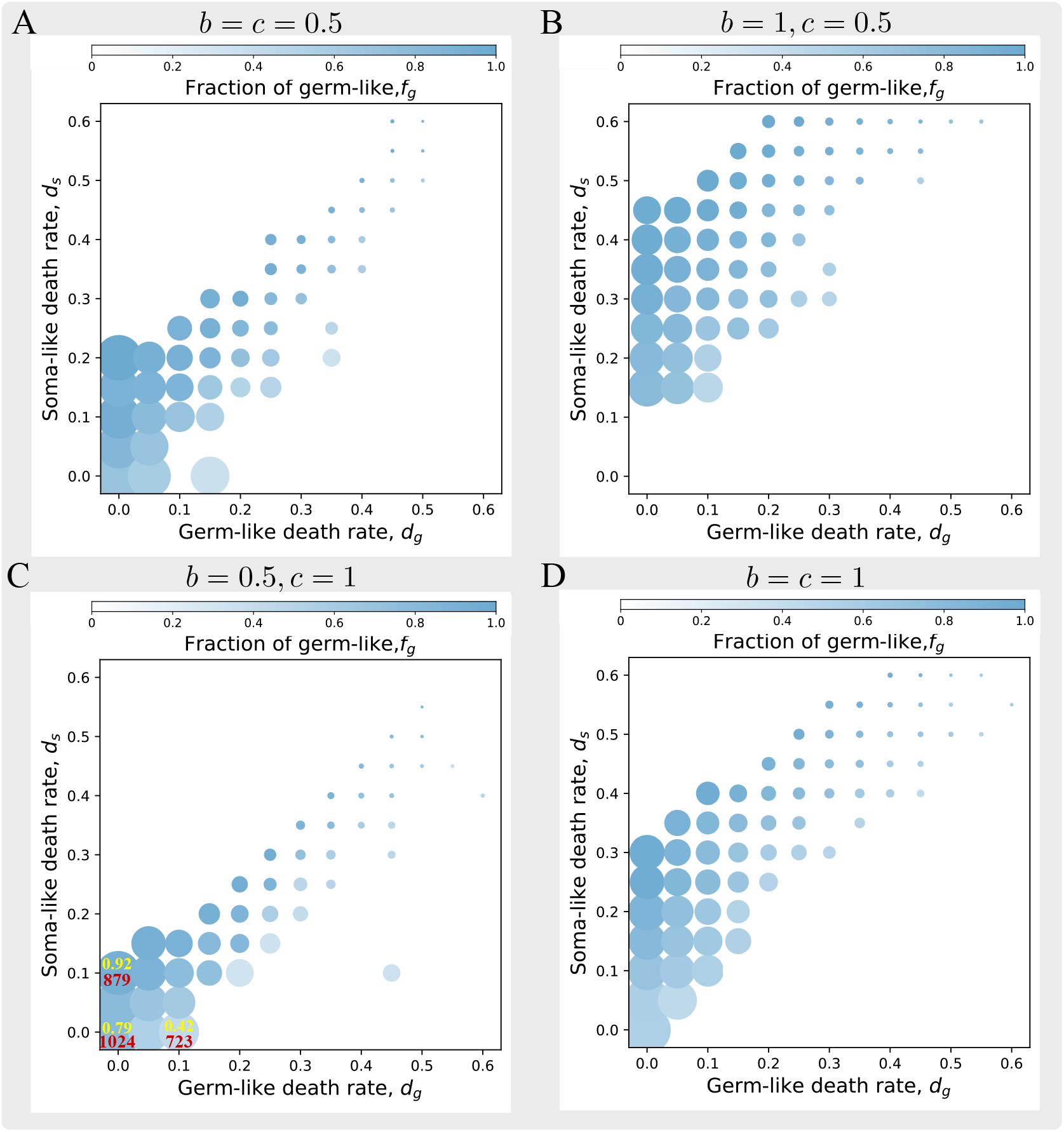
Cell death rates decrease cell number and cell fraction in organisms with *ID*. The averaged cell number and cell proportion of the organisms with *ID* strategy under *b* = 0.5, *c* = 0.5 in panel **A**, *b* = 1, *c* = 0.5 in panel **B**, *b* = 0.5, *c* = 1 in panel **C** and *b* = 1, *c* = 1 in panel **D**.The bubble size represents the average cell number of the maturity organisms with *ID* strategy. The bubble color represents the averaged proportion of germ-like cells within the organisms with *ID* strategy. The red and yellow numbers in panel **C** represent the specific values of the total number of cells and the proportion of germ-like cells, respectively. Parameters: *n* = 10, *α* = 1. At each pixel, 10 replicates for calculating the optimal strategy.

**Figure 5.**
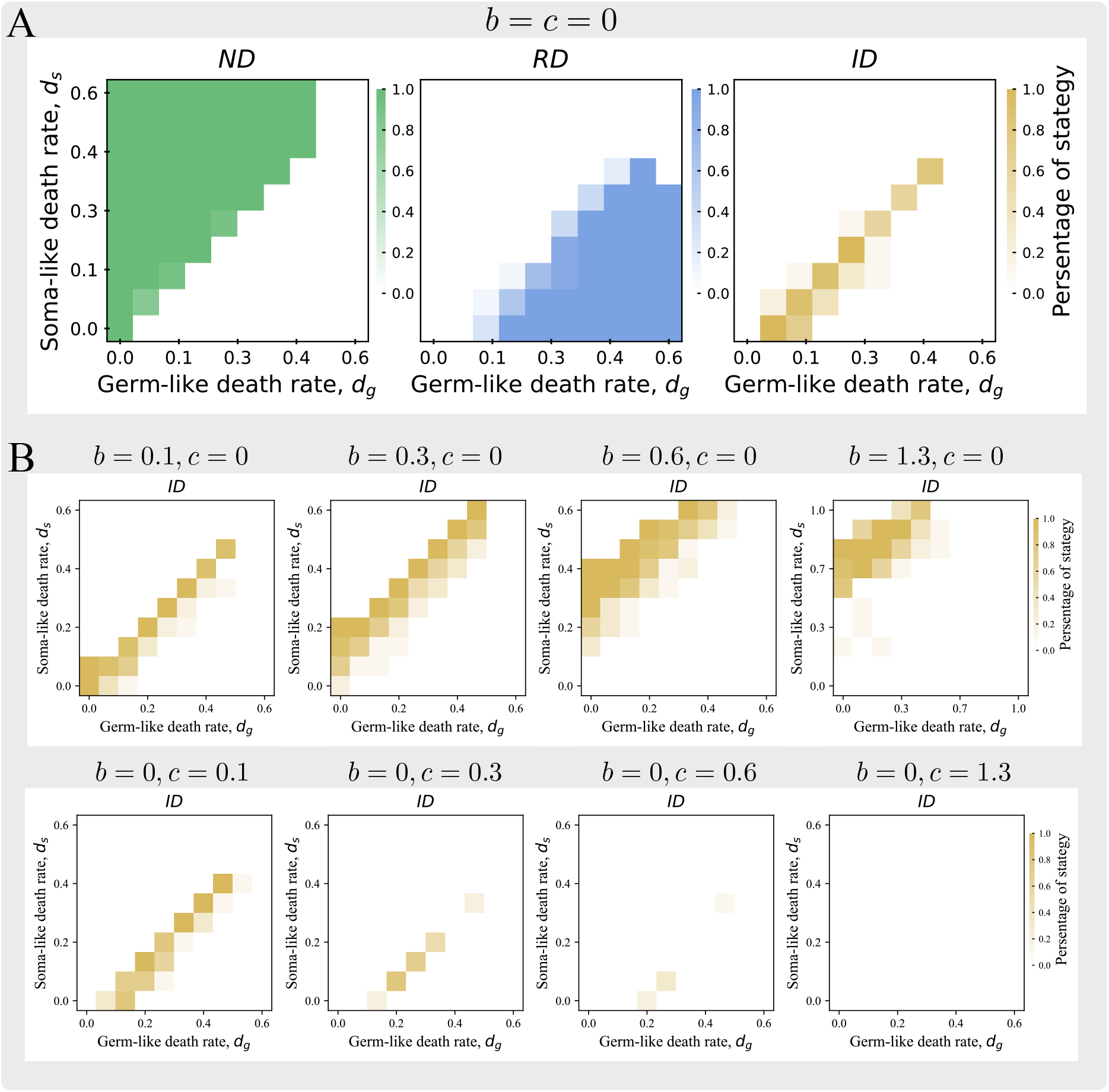
*ID* emerges mostly when *d*_*g*_ and *d*_*s*_ show linear relationship. A. The percentages of *ND, RD* and *ID* being optimal under varying cell death rates with no differentiation benefits and costs i.e. *b* = 0, *c* = 0. **B**. The percentages of *ND, RD* and *ID* being optimal under varying cell death rates, differentiation benefits and costs. Other parameters: *n* = 10 and *α* = 1. Replicates for searching optimal strategy at each dot is 10.

## 4 Conclusion and discussion

In the paper, we adopted a multicellular organism model to investigate the impact of cell death on the evolution of irreversible cell differentiation by considering cell interaction effects of differentiation benefits and costs. Irreversible differentiation describes the loss process of cell differentiation ability and is defined by 0 differentiation probability at the last cell division [Gao et al., 2023]. Our results show irreversible differentiation emergencies mostly when the cell death rates of germ-like cells and soma-like cells change in a linear relationship. The result indicates that the cell death difference between two closely related cell lineages shouldn’t be too large to evolve irreversible differentiation. It also suggests regulated cell death in an organism needs to keep a close cell death on different cell types. Then, we further found that irreversible differentiation can occur in new parameter space by implementing different cell death rates on cell types. Specifically, our results show that irreversible differentiation extends to high differentiation costs areas when germ-like cells die (Fig 3). Meanwhile, the emergence condition of irreversible differentiation is not influenced by equal cell death rates. The result implies the factors that cause the same death in different cell types do not change the organism’s differentiation patterns. Finally, both cell death rates can reduce the cell number and the fraction of their cell types in mature organisms. However, we found that germ-like cell death has more stronger impact on reducing organismal size and cell fraction compared with soma-like cell death.

In the model, we assumed the two cell types are based on their life functions as the precise definition of cell type is still lacking, instead functional and morphological differences is viewed as characteristics of distinct cells [levers et al., 2017, Márquez-Zacarías et al., 2021]. The two cell types are more like general cells but with different task preferences rather than specialized cell types. In other words, we investigated the cells that have functional preferences but are still in the state before finally turning into specialized cells by irreversible differentiation. These transient cell types allow us to investigate the cell differentiation patterns which need to screen the differentiation probabilities along with cell divisions. A similar way of defining cell types by cells’ function in other theoretical investigations of cell differentiation [Michod, 2007, Rueffler et al., 2012, Ispolatov et al., 2012, Rodrigues et al., 2012, Yanni et al., 2020, Cooper et al., 2022]. An alternative way of cell types definition is the Boolean model, which is a discrete dynamical system comprising a set of Boolean variables (e.g., True/False, 0/1), the values depending on a set of functions applied to each variable [Márquez-Zacarías et al., 2021]. Another method of cell types definition is via the fixed point attractors of a continuous dynamical system which is controlled by the expression levels of related genes [Nakamura et al., 2024].

One of the differences between our work and other theoretical investigations of cell differentiation is the differentiation pattern. Previous work mostly focuses on the differentiation state of cells rather than the differentiation patterns which are the differentiation trajectories of cells [Michod, 2007, Rueffler et al., 2012, Ispolatov et al., 2012, Yanni et al., 2020, Cooper et al., 2022]. Differentiation patterns of irreversible and reversible have been considered in filament-shaped organisms with two indispensable cell types [Rodrigues et al., 2012]. Comparatively, our model contains organisms born with single germ cells and meanwhile can include the strategy of no differentiation strategy. Another difference is the fitness definition of organisms. To quantify cell differentiation, many models adopted the product of the tasks undertaken by different cells [Michod, 2007, Rueffler et al., 2012, Ispolatov et al., 2012, Yanni et al., 2020, Cooper et al., 2022]. However, we use an organism’s reproductive rate as a fitness proxy to evaluate differentiation strategies. One of the reasons is that cell number and composition continuously change with cell death effects. Thus, the product of the tasks split by cells is hard to quantify as it continuously changes. Another reason is that differentiation patterns like irreversible differentiation need to consider cells’ differentiation probabilities that accompanied with cell division series. Thus, we took the final reproductive rate as a fitness proxy.

Additionally, we investigated the irreversibility of cells under the cell division times *n* = 10. We chose this value because *Volvox*, the model-inspired organisms, contains thousands of cells in total [Matt and Umen, 2016]. Thus by taking *n* = 10, an organism will grow to contain 1024 cells in total without cell death. Altogether, our model provides an analytical method to investigate different types of cell death effects on organisms’ differentiation patterns. In the model, the cell death rate is designed as a stage-dependent variable 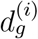 and 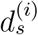. Although we take constant cell death rates *d*_*g*_ and *d*_*s*_ for convenience, cell death can be designed as a pair of functions depending on the factors causing cell death. For example, cell death is highly likely to happen for some cells rather than others under the same environmental stimulus [Ratcliff et al., 2012]. Aside from small multicellular organisms, our model also provides a model for cell differentiation patterns in tissues. The model can also be extended to include more cell types and thus a hierarchy of differentiation patterns.

## Data availability

Numerical data produced by this work has been deposited in GitHub: https://github.com/YuanxiaoGao/Thevolution-of-irreversible-cell-differentiation-under-cell-death.

## A Appendix

### A.1 Cell number of maturity organisms under cell death

With cell death, the cell number of a maturity organism *N* ^(*n*)^ changes with varying cell death rates. In our study, we considered two types of cell death scenarios: (i) the unit cell death rates of germ-like and soma-like cells were the same at each division i.e. 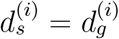, and (ii) the unit cell death rates of germ-like and soma-like cells were different at each division i.e. 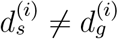. Next, provide a detailed description.

#### Cell number of a maturity organism *N* ^(*n*)^ **under** 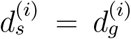

If the unit cell death rate of germ-like cells and soma-like cells is the same, then we assume 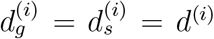. Assuming the total number of cells after the *i*th division is *N* ^(*i*)^, the total number of cells after undergoing a synchronous division is 2*N* ^(*i*)^, and the time consumed is *t*^(*i*+1)^. During this period, the total number of dead cells was 2*d*^(*i*)^*N* ^(*i*)^*t*^(*i*+1)^. So the total number of cells *N* ^(*i*+1)^ during the *i* + 1st division is expressed as the difference between the total number of cell death not considered and the dead cells, and is denoted as:

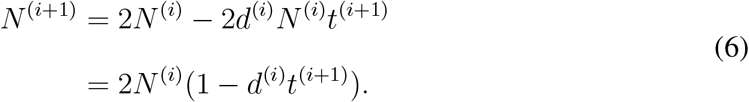

By recursion, the total number of cells *N* ^(*n*)^ during the last division is

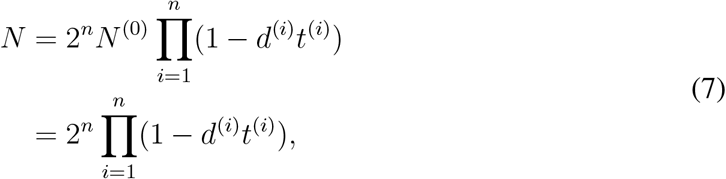

where *N*^(*i*)^ represents the total number of cells after the *i*th division, 2*d*^(*i*)^*N* ^(*i*)^*t*^(*i*+1)^ represents the total number of cells that died within the time *t*^(*i*+1)^ spent on the *i* + 1th division, and *N* ^(0)^ represents the total number of cells before the first division. Since cell division begins with a single germ-like cell, *N* ^(0)^ = 1.

#### Cell number of a maturity organism *N*^(*n*)^ **under** 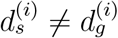

If the unit cell death rates of germ-like cells and soma-like cells are different, then the number of germ-like cells and soma-like cells are two independent quantities at each division. We assume that the total number of germ-like cells is 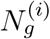and the total number of soma-like cells is 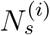, where *I* represents the *i*th division and

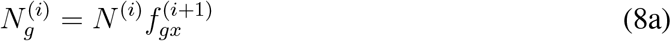

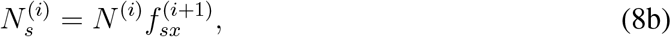

According to the calculation method of (i), we provide the total number of cells at the (*i* + 1)th division *N* ^(*i*+1)^,

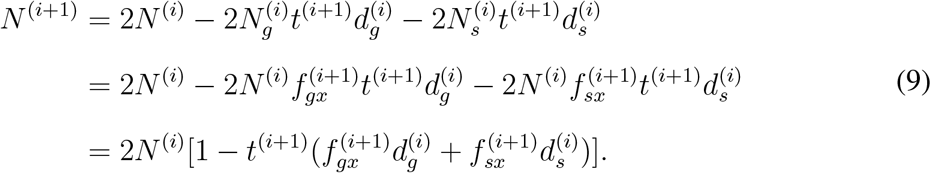

By recursion, the total number of cells *N*^(*n*)^ during the last division is

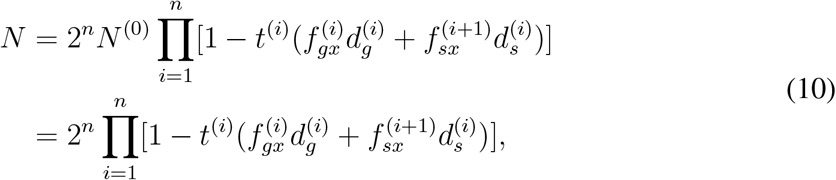

where 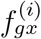 represents the proportion of germ-like cells in the *i*th division, 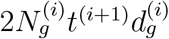 represents the total number of all germ-like cells that died within the time *t*^(*i*+1)^ spent on the *i* + 1th division, and 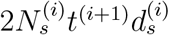 represents the total number of all soma-like cells that died within the time *t*^(*i*+1)^ spent on the *i* + 1 division.

### A.2 Numerical calculation of searching for the optimal strategy

#### Differentiation strategy space

We the same numerical method to calculate an organism’s reproductive rate [Gao et al., 2023]. In the model, the differentiation strategies are infinite. To simplify, we use discrete differentiation probabilities to capture the effects of all differentiation strategies. We use grid search to find all possible differentiation strategies. Thus, we set *δ*^(*i*)^ = 0.05 or *δ*^(*i*)^ = 0. Then, differentiation probabilities 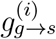 and 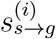 are confined in the values of 0, 0.05, 0.1, …, 1. That is division probabilities 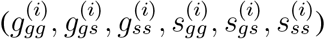 are confined in the values of 0, 0.1, 0.2, …, 1. Then, we receive the first set of differentiation probabilities of a strategy 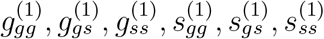. Then, with a difference of 0.1 or 0 randomly generate the sets of differentiation probabilities for each cell division until *n* times. Even with the discrete differentiation space, there are also numerous differentiation strategies. For example, if *n* = 10, we have 4356 choice for the first cell division, then each of them will have at least (for boundary conditions) 5 sets and at most 13 sets of differentiation probabilities for the second cell division. That is we will have at least 4356 *×* (2^5^)^10^ strategies in the generated differentiation probability space. We cannot explore all the strategies and calculate their reproductive rates for comparison. Instead, we use the Monte-Carlo methods randomly to sample differentiation strategies and then calculate and compare the reproductive rates of organisms under these strategies.

#### Search for the optimal differentiation strategy

To find the optimal strategy at a fixed parameter space of cell death, benefit, and cost, we first randomly choose 100 number of the first cell differentiation probabilities in the 4356 strategy space, then for each of them, we randomly choose 10 strategies. Then there are 1000 strategies in total. Then we calculate the reproductive rates of these strategies and find the strategy with the largest reproductive rate. Considering the huge number of strategies, we run the above process several duplicates depending on the circumstances. Then we get the percentages of each classified strategy being optimal at the fixed parameter values of cell death, benefit, and cost. Finally, we repeat the above calculation across all possible parameter spaces in terms of cell death, benefit, and cost.

### A.3 Irreversible differentiation emerges mostly under linear cell death rates of different cell types

The results show that irreversible differentiation emerges mostly when *d*_*g*_ and *d*_*s*_ show a linear relationship under different cell interactive parameter spaces. Irreversible differentiation emerges at a higher soma-like death rate when differentiation benefits increase. In comparison, irreversible differentiation decreases when differentiation costs increase.

